# Paternal hypoxia exposure primes offspring for increased hypoxia resistance

**DOI:** 10.1101/2020.12.09.416727

**Authors:** Alexandria Ragsdale, Oscar Ortega-Recalde, Ludovic Dutoit, Anne A. Besson, Jolyn H.Z. Chia, Tania King, Shinichi Nakagawa, Anthony Hickey, Neil J. Gemmell, Timothy Hore, Sheri L. Johnson

**Affiliations:** Department of Zoology, University of Otago, Dunedin, New Zealand; Department of Anatomy, University of Otago, Dunedin, New Zealand; Evolution and Ecology Research Centre, School of Biological, Earth and Environmental Sciences, University of New South Wales, Sydney, NSW, Australia; School of Biological Sciences, University of Auckland, Auckland, New Zealand

## Abstract

In a time of rapid environmental change, understanding how the challenges experienced by one generation can influence the fitness of future generations is critically needed. Using tolerance assays, transcriptomic and methylome approaches, we use zebrafish as a model to investigate transgenerational acclimation to hypoxia. We show that short-term paternal exposure to hypoxia endows offspring with greater tolerance to acute hypoxia. We detected two hemoglobin genes that are significantly upregulated by more than 7-fold in the offspring of hypoxia exposed males. Moreover, the offspring which maintained equilibrium the longest showed greatest upregulation in hemoglobin expression. We did not detect differential methylation at any of the differentially expressed genes, suggesting that another epigenetic mechanism is responsible for alterations in gene expression. Overall, our findings suggest that a ‘memory’ of past hypoxia exposure is maintained and that this environmentally induced information is transferred to subsequent generations, pre-acclimating progeny to cope with hypoxic conditions.

## INTRODUCTION

Paradigm-shifting research has revealed that the life-history experiences of parents can influence the phenotype of their offspring through non-genetic mechanisms (Salinas et al., 2013; Bohacek and Mansuy, 2015; O’Dea et al., 2016; Dias and Ressler, 2013; Gapp et al., 2014; Radford et al., 2014; Burton and Metcalfe, 2014; Bonduriansky, 2012; Bonduriansky and Day, 2008). Non-genetically transmitted phenotypes can be generated by diverse environmental effects, affecting a wide array of offspring traits, both positively and negatively (O’Dea et al., 2016). Work on mice has demonstrated that learned fear responses (Dias and Ressler, 2013; Gapp et al., 2014) and metabolic alterations associated with undernourishment can be inherited via sperm (Radford et al., 2014). These studies suggest that parents can transmit information that may benefit offspring survival. Through this transgenerational plasticity (also known as transgenerational acclimation (Herman and Sultan, 2011; Marshall, 2008)), parents may provide offspring with increased tolerance to environmental perturbations, such as contaminants (Araujo et al., 2019; Kishimoto et al., 2017; Marshall, 2008), food shortages (Kishimoto et al., 2017; Weyrich et al., 2018), carbon dioxide (Allan et al., 2014; Shi et al., 2020; Lee et al., 2020), hypoxia (Ho and Burggren, 2012), but see (Truebano et al., 2018), salinity (Heckwolf et al., 2020, 2018), and temperature (Donelson et al., 2014, 2012; Veilleux et al., 2015; Salinas and Munch, 2012; Weyrich et al., 2018; Ryu et al., 2018). Studies have tended to focus on maternal transgenerational plasticity, or have exposed both parents to the environmental perturbation, making it impossible to disentangle the relative roles of mothers and fathers in altering offspring phenotype (Guillaume et al., 2016; Rutkowska et al., 2020). Thus, a better understanding of the specific role of paternal effects in transgenerational plasticity is needed, especially considering environmental specific information is likely transferred via sperm (see below). Further, the underlying molecular processes have been identified in a just a few studies (Heckwolf et al., 2020; Kishimoto et al., 2017; Ryu et al., 2018; Shi et al., 2020; Strader et al., 2019; Veilleux et al., 2015). For example, metabolic genes are upregulated transgenerationally in the damselfish (*Acanthochromis polyacanthus*), suggesting shifts in energy production for maintaining performance at elevated temperatures (Veilleux et al., 2015). Potential epigenetic mechanisms for this transgenerational acclimation have been detected, via differential methylation of genes involved in energy homeostasis, mitochondrial activity, and oxygen consumption (Ryu et al., 2018). In a time of rapid environmental change, a better understanding of how environmental challenges experienced by organisms could increase the fitness of future generations to survive these same stressors is critically needed.

Hypoxia, defined as sufficiently low levels of oxygen to deprive tissues of oxygen, is a major physiological challenge (Diaz and Rosenberg, 2008; Long et al., 2015; Wang et al., 2016). The aerobic lifestyle of most animals requires a constant supply of sufficient oxygen, and low oxygen levels constitute a major environmental threat (Roesner et al., 2006). Hypoxic conditions precipitate conserved physiological effects in a wide array of vertebrates (Okumura et al., 2003; Saxena, 1995; Wu et al., 2003), but hypoxia is particularly well-studied in aquatic species. The total oxygen present in water at 30°C is only 0.5% that of air; hence, physiological stress of hypoxia challenges many freshwater and marine organisms, and hypoxia is one of the most widespread issues in aquatic habitats due to the rise of dead zones and climate change (Diaz and Rosenberg, 2008; Jenny et al., 2016). Thus, it is now more important than ever to understand how different aquatic species will response to environmental hypoxia.

The negative impacts of hypoxia on growth and reproduction are well-recognized (Townhill et al., 2017; Wu, 2002; Wu et al., 2003), with recent studies also discovering transgenerational effects, whereby marine medaka fish (*Oryzias melastigma*) exposed to hypoxia show reproductive impairments in F1 and F2 generations, suggesting that hypoxia might pose a long-lasting threat to fish populations (Lai et al., 2019; Wang et al., 2016). But fishes, in particular, are also notable for their adaptive abilities to acclimate to hypoxic conditions (Alexander et al., 2017; Diaz and Rosenberg, 2008; Nikinmaa, 2002; Richards, 2011), including physiological, morphological, and phenotypic responses. Remarkably, goldfish (*Carassius auratus*) and crucian carp (*Carassius carassius*) exhibit gill remodeling in response to hypoxia exposure (Dhillon et al., 2013; Tzaneva et al., 2011). Zebrafish embryos and larvae modify cardiac activity and blood vessel formation in response to hypoxic exposures (Pelster, 2002). In addition to morphological alternations, gene expression can also be affected by hypoxic conditions. In zebrafish gill tissue, more than 300 genes are differentially expressed between hypoxia exposed individuals versus controls; these changes in gene expression are coupled with morphological changes in gill structure, such as increased surface area of gill tissue (van der Meer et al., 2005). Prolonged exposure to low oxygen has been shown to improve hypoxia tolerance in Murray cod (*Maccullochella peelii*; (Gilmore et al., 2020) and snapper (*Pagrus auratus;(Cook* et al., 2013)). Likewise, pre-acclimation of zebrafish larvae to mild hypoxia significantly improves their resistance to lethal hypoxia, and upregulation of some oxygen transport genes are associated with this acclimation (Long et al., 2015). Intriguingly, zebrafish offspring of males and females exposed to chronic hypoxia have a higher resistance to acute hypoxia than those of controls (Ho and Burggren, 2012), despite the offspring having never been exposed to hypoxia, suggesting that parents may pass on information that may pre-acclimate offspring to cope with hypoxic conditions.

Gene expression and transgenerational effects can be regulated by epigenetic modifications, including DNA methylation, histone modifications, and non-coding microRNAs (O’Dea et al., 2016). For example, in the marine medaka studies, transgenerational effects are associated with differential methylation in sperm and ovary, with altered gene expression in genes known to be associated with spermatogenesis and gene silencing (Wang et al., 2016) and cell cycle control and cell apoptosis (Lai et al., 2019). DNA methylation has also been investigated as a means underlying transgenerational effects of chemical exposures in zebrafish (Carvan et al., 2017) and the paternal methylome is believed to be stably transmitted to offspring in zebrafish (Jiang et al., 2013; Potok et al., 2013) without global reprogramming in primoridal germ cells (Ortega-Recalde et al., 2019; Skvortsova et al., 2019), potentially facilitating environmental specific information transfer.

Here we use zebrafish to further explore the phenomenon of transgenerational acclimation of hypoxia in fishes. We focus on paternal exposure as we predict that environmental specific information can be transferred via sperm, as observed in other studies (Dias and Ressler, 2013; Lamb et al., 2020; Wang et al., 2016). We test whether paternal exposure to hypoxia stimulates phenotypic responses in offspring, using behavioural phenotyping to identify resistance to acute hypoxia (time to loss of equilibrium). We then use RNA-Seq to identify candidate genes that are differentially expressed in control and hypoxic progeny, and whole genome bisulfite sequencing data to assess whether changes in DNA methylation underpin alterations in phenotype and gene expression (Figure 1).

**Figure 1.**
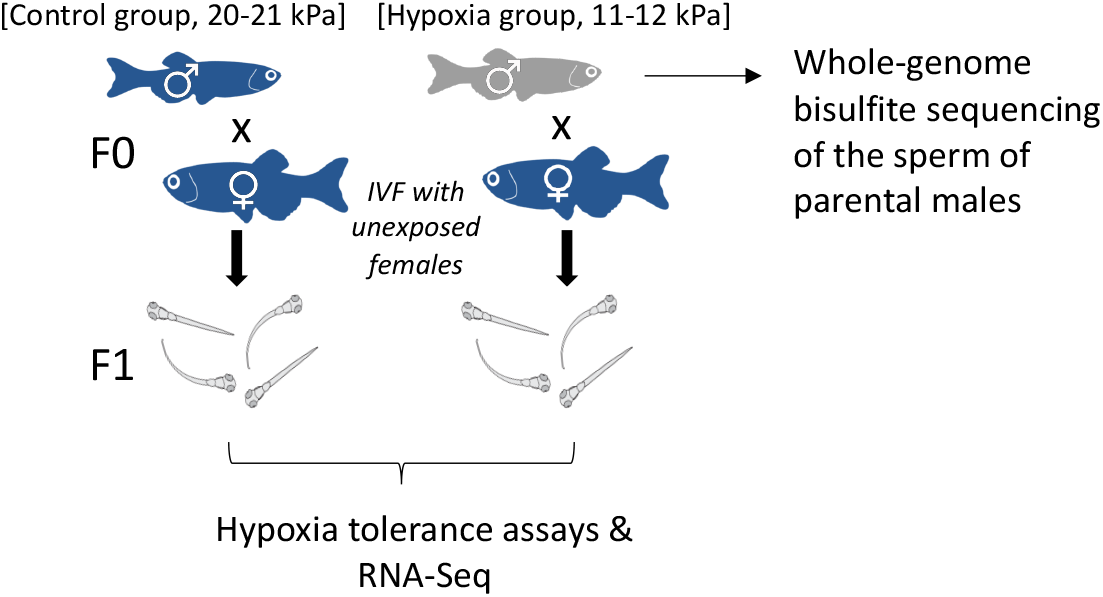
Experimental design. Groups of adult male zebrafish are maintained in normoxia (20-21 kPa, n = 20) or hypoxia (11-12 kPa, n = 20) for 14 days. Five males from each treatment were then used to create F1 progeny, crossing the males to unexposed females (n = 5 families/treatment), with half of the sperm used for whole genome bisulfite sequencing, to assess differential methylation (n = 3 per treatment). At 20-21 days post fertilization, offspring undergo acute hypoxia tolerance (6-8 offspring/family) assays and n = 3 offspring/treatment are used for differential gene expression analysis (RNA-Seq).

## MATERIALS AND METHODS

### Fish husbandry

Breeding and husbandry took place within the Otago Zebrafish Facility (OZF), a temperature-controlled facility maintained at 25-27°C, pH 7-7.8 and conductivity 300-500 μS. Fish were maintained in a Tecniplast re-circulating system (Tecniplast, Varese, Italy) under a 14:10 light:dark photoperiodic cycle, with 30 minutes of simulated dawn and dusk at the start and end of each day. Acute hypoxia assays took place in the Zoology Department, where room temperature was controlled at 25°C, with a 13.5 hr (0700-2030 hr) light cycle with 30 min of simulated dawn and dusk. Zebrafish were fed twice daily with dry food (ZM000-400, size-dependent) and once daily with live rotifer (*Brachionus* spp. days 5-10 post fertilisation) or artemia (*Artemia salina*; days 10+ post fertilization). All animals were collected and maintained according to the standards of the Animal Ethics Committee for the University of Otago, New Zealand (protocol no. AEC 44/16).

### Hypoxia exposure

In November 2016, nine-month-old male zebrafish (AB wild-type; n=20/treatment) were exposed to hypoxic conditions (10.91-12.33 kPa *p*O_2_) or control conditions (20-21 kPa), for two weeks. Two glass tanks (36×29×26.5 cm) were separated into three zones (12 x 29 cm), with 10 fish in each outer compartment and two fine bubble diffusers and a filter positioned in the middle compartment. The oxygen concentration of the treatment tank was maintained by using an OxyGuard Mini probe (OxyGuard International, Denmark) and oxygen controller (Model PR5714, PR Electronics, Denmark) that were connected to nitrogen and air cylinders (BOC Gas Supplies, Food Fresh grade). An on/off relay output from the controller actuated a solenoid-controlled flow of compressed nitrogen or compressed air (BOC Gas Supplies, New Zealand), to maintain the system at 53.1-60 % air saturation (= 10.91-12.33 kPa or 4.38-4.95 mg/L) level (Cook et al., 2013). Normoxic conditions in the control tank (>95% saturation) were maintained by continually passing air through the bubble diffusers, connected to an air pump. The oxygen concentration in the tanks was confirmed with a YSI 85 probe (YSI, Inc., Ohio, USA). Both tanks were siphoned for waste and received 10% water changes every three days. Calibration of the oxygen probe was checked daily and recalibrated as necessary.

### Hypoxia tolerance assays and analysis

Progeny from five families per treatment (n=5 hypoxia, n=5 control) were generated by *in vitro* fertilisation (IVF) (Johnson Sheri L. et al., 2018; Lamb et al., 2020) seven days after parental exposure finished. Briefly, eggs and sperm were collected using abdominal massage – half the sperm was used for IVF and the other half was stored for subsequent DNA extraction. Offspring were reared to 20-21 dpf when n=8 offspring per family (except family C5, where n = 6; total n = 77) were challenged by acute hypoxia assays in small acrylic chambers (100 mm H x 50 mm x 50 mm). A 0-1 kPa oxygen level was achieved by continually passing compressed nitrogen through a bubble diffuser for at least 10 minutes before assays and continuously passing the nitrogen through a valve in the top of the chambers during the assay. The oxygen concentration fluctuated between 0 and 1%, monitored using a fibre-optic oxygen probe (Foxy OR-125) attached to an Ocean Optics^®^ USB 2000 spectrophotometer with USB-LS-450 light source and the manufacturer’s software (OOI Sensors). Each fish was filmed for 240 seconds and recorded with a GoPro Hero 3+ camera. A total of 77 videos (n=37 control, n=40 hypoxia) were tracked using EthoVision XT behavioral tracking software, version 11.5 (Noldus Information Technology, Netherlands). Fish were then immediately euthanized by submersion in ice and stored in RNAlater (Invitrogen). One offspring from family C1 stayed at the bottom the entire assay, so this individual was removed from further analyses.

Resistance to acute hypoxia, defined as the first time that progeny lost equilibrium for 3 seconds or more during hypoxia assays (Ho and Burggren, 2012), and the loss of equilibrium frequency were analysed using R v. 3.5.1 (R Core Team, 2018). Loss of equilibrium > 3 s was initially modelled using a gaussian generalized linear model, and loss of equilibrium frequency as a poisson generalized linear model, with treatment (control vs. hypoxia) as a fixed effect. To account for multiple fry per family being tested, we also ran the models incorporating Treatment ID as a random effect, using a gaussian linear model for time to loss of equilibrium > 3 s and a poisson linear mixed effects model for the loss of equilibrium frequency. Significant heterogeneity was obvious in the time to loss of equilibrium, so a linear mixed model incorporating heteroscedasity was compared to the model assuming homogeneity, by incorporating the variance in Treatment into the model (Cleasby and Nakagawa, 2011).

### RNA-Seq library preparation and analysis

Total RNA from six whole 20-21 dpf offspring (3 control offspring, 3 treatment offspring) was extracted using a Zymo Duet extraction kit (Zymo, NZ). The integrity of RNA samples was determined using an Agilent RNA 6000 Nano chip on an Agilent 2100 Bioanalyzer to check that the samples had an RNA Integrity Number (RIN) value of 8–9. Total RNA concentration was measured by Qubit 2.0 Fluorometer (Qubit RNA HS Assay Kit, Life Technologies). Samples were sent to the Otago Genomics and Bioinformatics Facility at the University of Otago, under contract to New Zealand Genomics Limited, for library construction and RNA sequencing. Messenger RNA sequencing libraries were prepared using the Illumina TruSeq Stranded mRNA sample preparation kit (Illumina), as per the manufacturer’s instructions. RNA sequencing was performed on the Illumina HiSeq2500 (Illumina, USA) machine with single-ended 100-bp reads generating 17.4-20.0 million reads per sample.

Low quality reads and remaining adapters were trimmed using Trim Galore v0.6.4 [https://www.bioinformatics.babraham.ac.uk/projects/trim_galore/] in a two-step process. First, sequencing adaptors were removed, 10 bp were hard-trimmed from the 5’ end to account for sequence bias produced by PBAT library preparation, and last, low-quality ends from reads (PHRED score <20) were removed (Bolger et al., 2014). The reads were then aligned to the zebrafish genome (GCRz11) using HISAT2 v2.2.0 (Kim et al., 2015), informed with the GCRz11.99 annotation. Expression was summarized sample by sample at the gene level using featureCounts v2.0.0 (Liao et al., 2014). The analysis of differential expression was conducted in R v.3.5.0 (R Core Team, 2018) using the DESeq2 package v1.24.0 (Love et al., 2014) without the independent filtering option implemented in the *results* function (see https://github.com/OscarOrt/Paternal_hypoxia_Ragsdale_2020). Differentially expressed genes between offspring of fathers exposed to hypoxia and controls were extracted after correcting for multiple testing using False Discovery Rate cut-off of q=0.05 (Benjamini and Hochberg, 1995a). In order to test for over/under-representation of biological pathways which differentially expressed genes are involved, enrichment of Gene Ontology (GO) analysis terms was performed using Gorilla online accessed on August 4 2020 (Eden et al., 2009).

### Whole genome bisulfite sequencing and analysis

DNA from six parental sperm samples was purified using a modified magnetic bead method (Peat et al., 2017). WGBS was undertaken using an adapted modified post-bisulfite adaptor tagging (PBAT) method (Miura et al., 2012; Peat et al., 2014). Briefly, bisulfite treatment was performed according to the EZ Methylation Direct Mag Prep kit (Zymo, D5044) instruction manual. Bisulfite treatment was performed before adaptor tagging, enabling simultaneous conversion of unmethylated cytosines, DNA fragmentation, and improving library preparation efficiency. Sequencing primers were added using random heptamer primers, and finally, sample-specific indexes and sequences required for Illumina flow-cells binding were added by PCR. Library integrity was assessed by agarose gel electrophoresis and a fragment analyser (Agilent) and sequenced on eight lanes (one flow cell) of an Illumina HiSeq using HiSeq2500 V4 sequencing of 100 bp single ended reads (in combination with 13 other samples, which were part of another study).

Raw reads were trimmed in Trim Galore v0.6.4 as previously mentioned. Read mapping was performed using Bismark v0.22.3 (Krueger and Andrews, 2011) with the option --pbat specified. Zebrafish genome version 11 (GRCz11) was used as reference. BAM files were deduplicated and reports containing methylation base calls were generated using the deduplicate_bismark and bismark_methylation_extractor scripts, respectively. The non-conversion rate during the bisulfite treatment was evaluated by calculating the proportion of non-CG methylation; by this measure, all libraries must have had a bisulfite conversion efficiency of at least 98.5% (Table S3).

CG methylation calls were analyzed in SeqMonk v1.47.0 [www.bioinformatics.babraham.ac.uk/projects/seqmonk]. To analyse methylation at gene level, probes with a minimum of 5 methylation calls were generated and the percentage methylation measured as number of methylated calls/total calls. CpG islands (CGIs) were identified using Gardiner-Garden & Frommer’s criteria and previously published datasets. For the former, 200-bp windows moving at 1-bp intervals were considered CGIs if the Obs/Exp value was greater than 0.6 and a GC content greater than 50%. For the latter, data obtained by biotinylated CxxC affinity purification (Bio-CAP) and massive parallel sequencing was used to identify non-methylated CpG islands (Long et al., 2013).

Coupling between methylation and gene expression in differentially expressed genes (DEG) and at genome level was interrogated using methylation levels at transcription start sites (TSS). TSS were defined as 200 bp centered on the first nucleotide of an annotated mRNA, and a threshold of at least 20 methylation calls to be included in the analysis. DEG were divided into underexpressed and overexpressed, whereas coupling at genome level was assessed dividing gene expression levels into quartiles. Custom annotation tracks were generated using Gviz v1.28.3 (Hahne and Ivanek, 2016).

## RESULTS

### Hypoxia tolerance assays in unexposed offspring

Progeny of males exposed to moderate hypoxia for 14 days show a greater resistance to acute hypoxia than progeny of control males – time to loss of equilibrium was, on average, 32 seconds longer for progeny of hypoxia exposed males (t = 2.52, p = 0.014; confidence interval (CI) = 7.38, 58.83; Figure 2A) and the progeny of hypoxia exposed males lost equilibrium, on average, 9.18 times vs. 12.32 times in the control progeny (t = −4.21, p <0.0001, CI = −0.433, −0.158; Figure 2B). However, when accounting for family (to avoid pseudo-replication due to multiple offspring per family being tested), the effect of time to loss of equilibrium becomes non-significant (t = 1.67, p = 0.133, CI = −12.66, 79.22; note this model accounts for observed heterogeneity in the data), though the effect remains for the number of times offspring lost equilibrium (z = −2.46, p = 0.014, CI = −0.567, −0.037). Hence, there are strong family effects (time to loss of equilibrium Treatment ID variance = 42.38; loss of equilibrium frequency Treatment ID variance = 0.025), with greater resistance to acute hypoxia being observed for just two families, H1 and H2 (Figure 2C and D).

**Figure 2.**
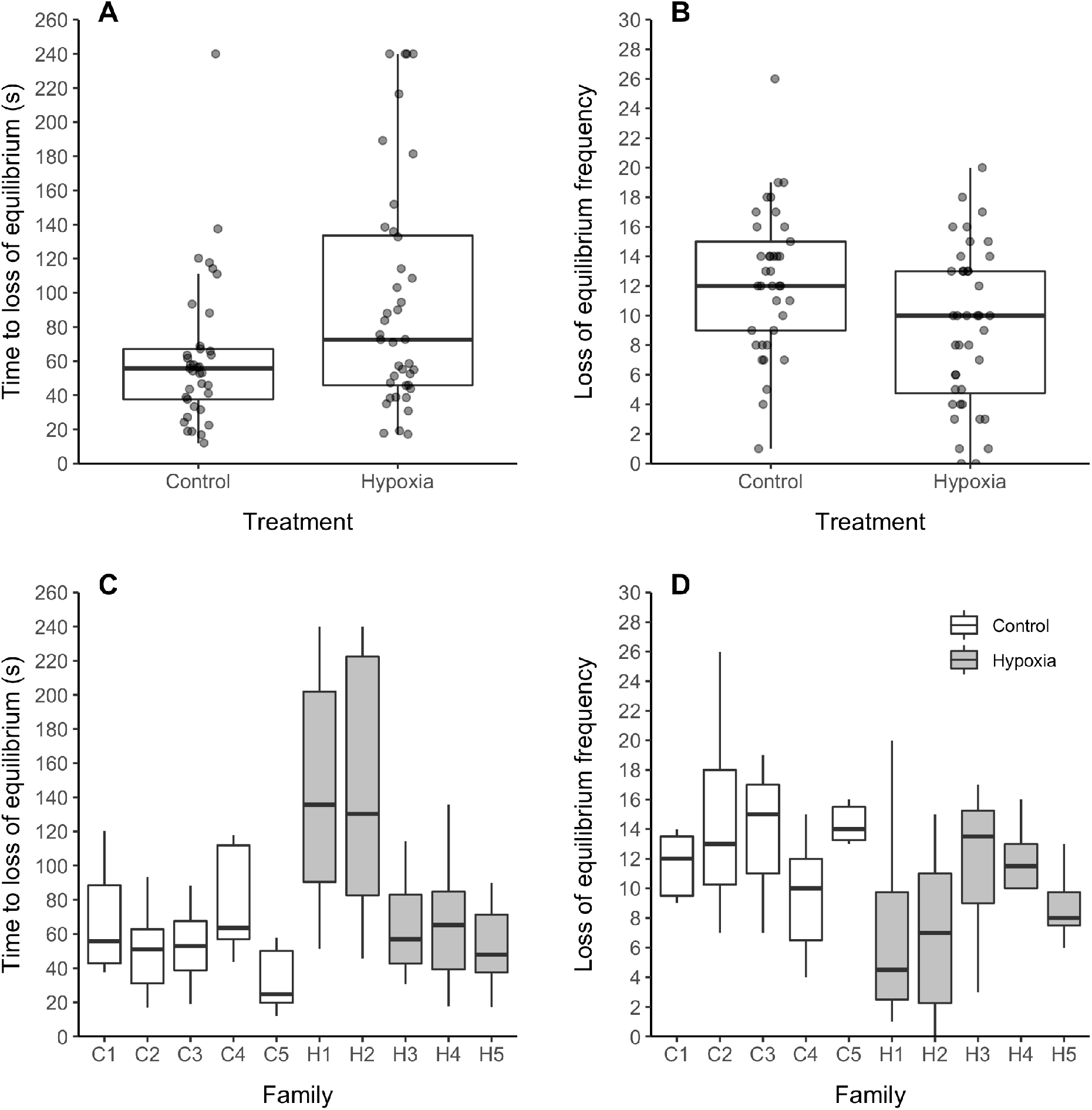
Hypoxia tolerance assays of unexposed offspring of control vs. hypoxia treated males. A and C) Time to first loss of equilibrium >3 sec, B and D) loss of equilibrium frequency – panels C and D are broken down by family. N = 6-8 progeny/family x 5 families/treatment. Tolerance assays done at 20-21 days post fertilization, following 14 day parental exposures (20-21 kPa normoxia control vs. 11-12 kPa hypoxia treatment). Boxes illustrate the interquartile range (medians, 25^th^ and 75^th^ percentiles), and whiskers illustrate 1.5 * the interquartile range, above and below the 75^th^ and 25^th^ percentiles.

### Transcriptome wide gene expression patterns in unexposed offspring

We sequenced 17.4 – 20.0 million reads per sample. Mapping assigned 14.1 – 16.1 million reads to genes, representing 80.15 – 81.41% of reads uniquely assigned to a gene, with detected expression in 26,260 of the 32,057 genes in the GRCz11.99 annotation.

A total of 91 genes were significantly differentially expressed between the offspring of control and hypoxia exposed males (Figure 3; Table S1). Of the 91 differentially expressed genes, 36 genes were significantly upregulated, and 55 were significantly downregulated in the offspring of hypoxia exposed males (Figure 3). Eight genes exhibited greater than 4-fold change in differential expression (2 upregulated in hypoxia, 6 downregulated in hypoxia) (Figure 4). Most notably, two hemoglobin genes (*hbaa1* and *hbz* (*si:ch211-5k11.6*); Figure 4 and 5A) were upregulated by more than 12-fold in offspring of hypoxia exposed F0 males. Three additional genes were upregulated by more than 7-fold, but did not pass False Discovery Rate (FDR) correction – these included a third hemoglobin gene (*hbba1*; Figure 4 and 5A; Table S2) and a major histocompatibility gene (*mhc1ula*; Figure 4). Importantly, the observed upregulation in hemoglobin gene expression correlates with the differential family effects observed in our loss of equilibrium tolerance assays (compare Figure 2 with Figure 5A/Table S2). Sequenced offspring from the H1 and H2 families, the two families that show the greatest tolerance to acute hypoxia, also show the greatest upregulation in gene expression. Indeed, the sequenced H1 offspring never lost equilibrium within our 240s long assay and the sequenced H2 offspring first lost equilibrium at 216.5s, whereas the third hypoxia offspring, from family H3 (with a first loss of equilibrium at 181.5s), and the offspring from C1-C3 (first loss of equilibrium > 3 s at 38.8, 24.2 and 18.9s, respectively) show little to no expression at these hemoglobin genes.

**Figure 3.**
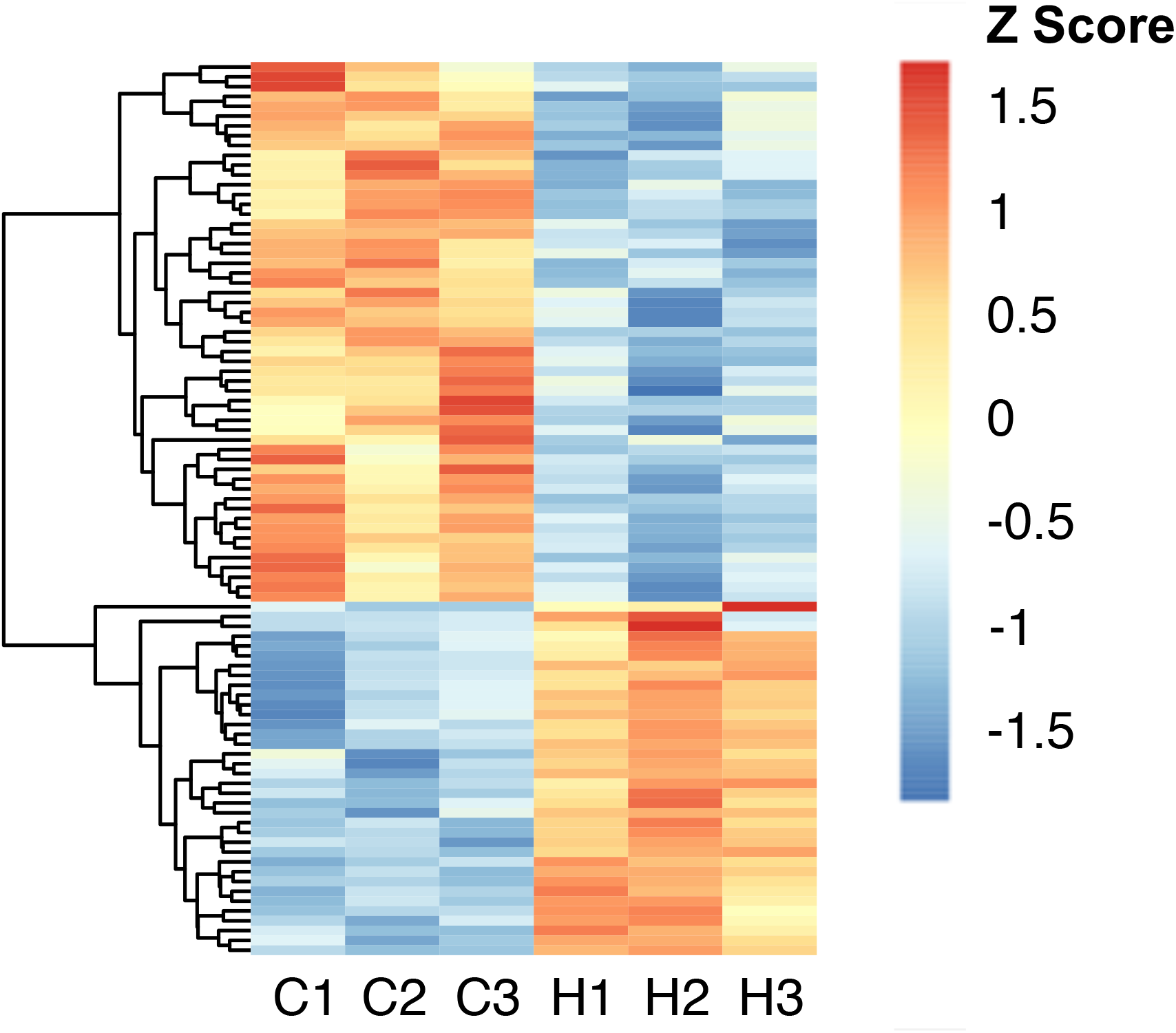
Differential gene expression in offspring of control vs. hypoxia treated males. 91 genes are differentially expressed between the 20-21 day old offspring of hypoxia (n = 3) and control (n = 3) males, with an FDR < 0.05. Red and blue colours represent higher and lower expression, respectively.

**Figure 4.**
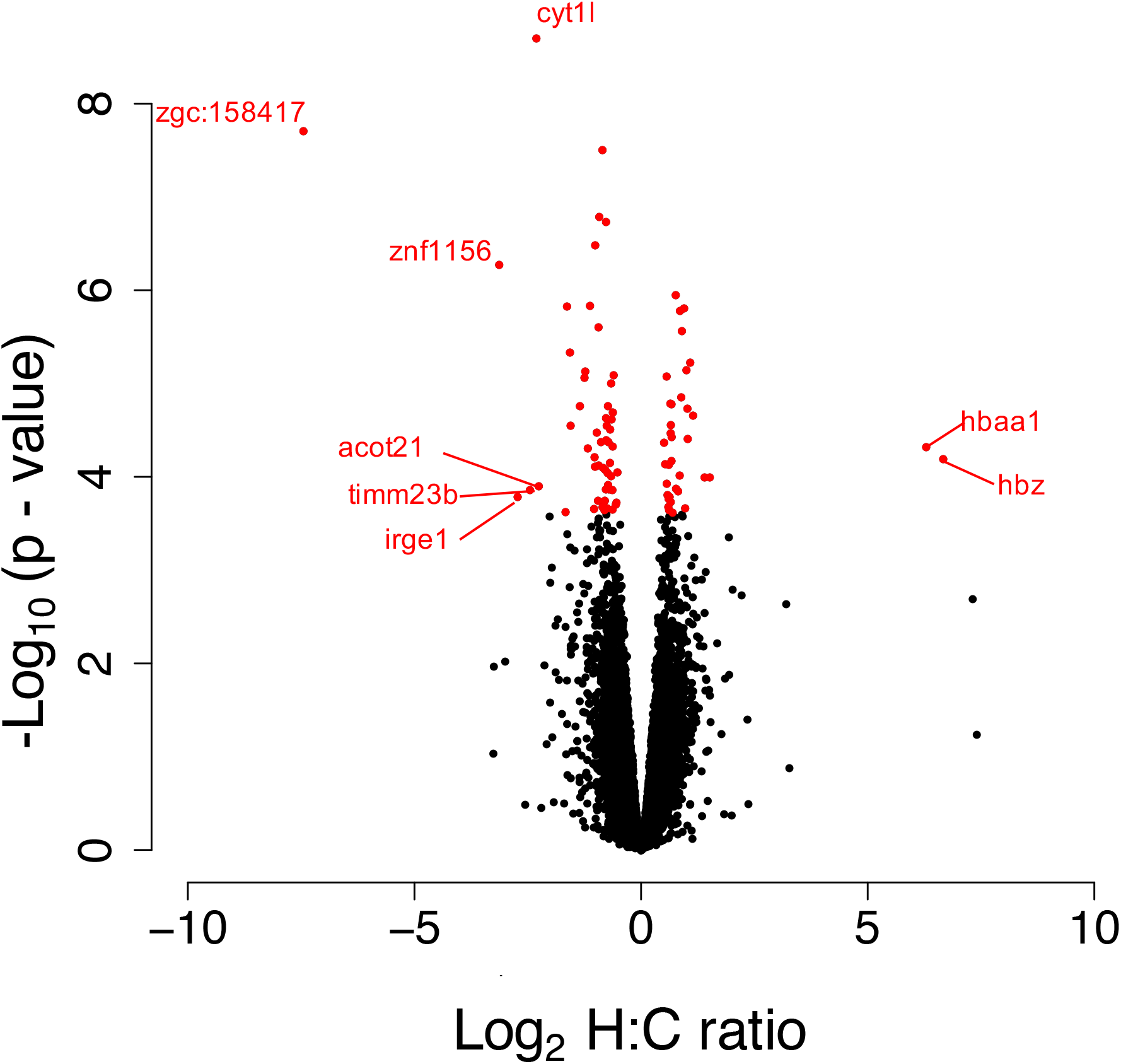
Volcano plot for differentially expressed genes in the offspring of control vs. hypoxia exposed males. 20-21 day old zebrafish F1 offspring from control (n = 3) and hypoxia (n = 3) treated males, showing the distribution of significance [-log10(p-value)] *vs*. fold change [log2(fold change)] for all genes. Each circle represents a gene, with significant genes (at 5% FDR) highlighted in red. Genes with greater than 4-fold change in expression between control and hypoxia treatments are labelled.

**Figure 5.**
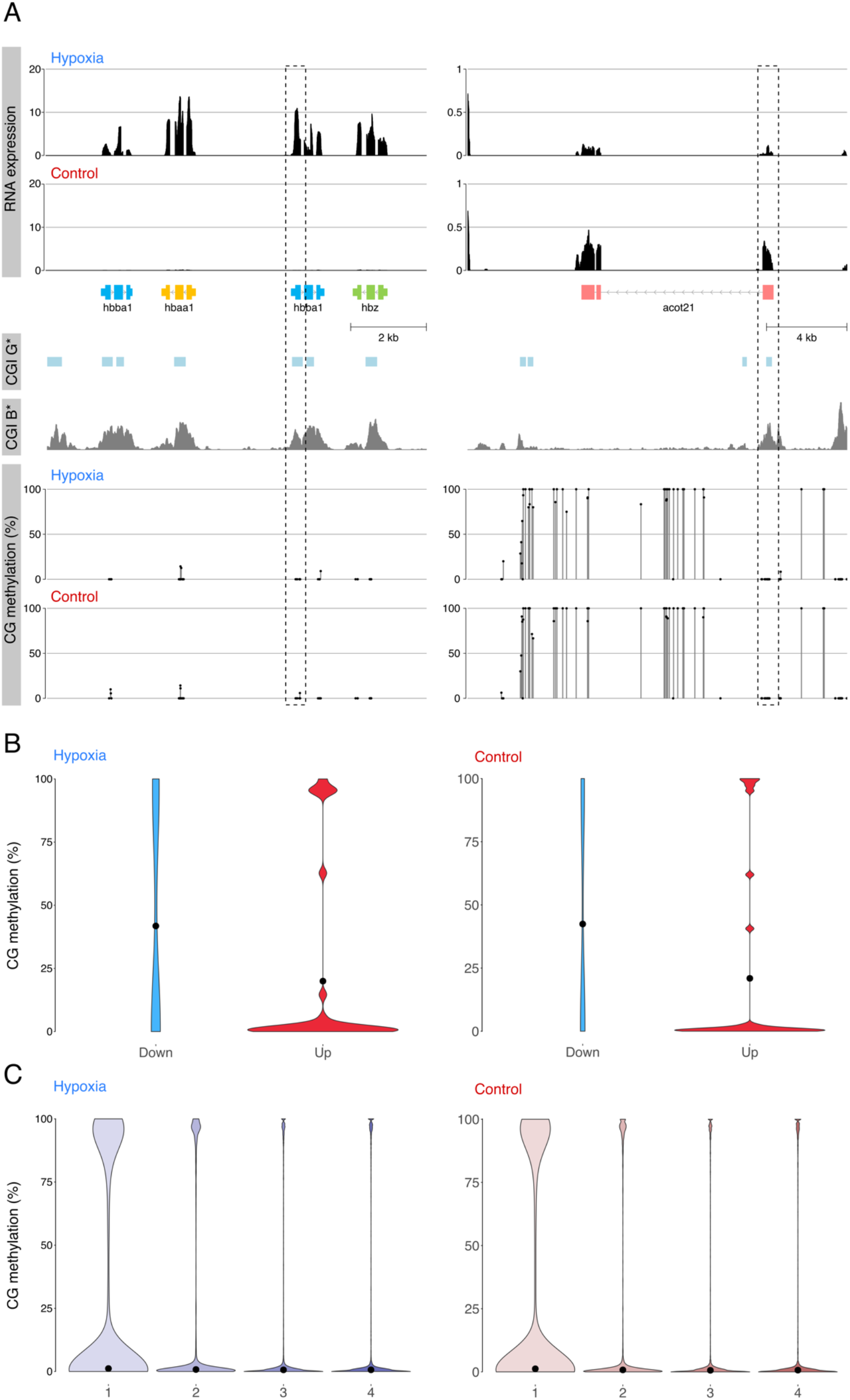
Global methylation changes and relationship between methylation in the sperm of hypoxia vs. control parental males and gene expression in the F1 progeny. A) Relationship between CpG methylation and RNA expression for Hb cluster and acot 21. CG methylation track shows methylation levels for dinucleotides with >5 calls; CGIs were predicted according to the Gardiner-Garden and Frommer criteria (CGI G*) and CxxC affinity purification (CGI B*). No changes in TSS methylation containing CGIs (dashed box) were observed for both genes (bottom). B) Violin plot showing distribution of methylation at transcription start sites (TSS) of differentially expressed genes in the F1 progeny. Downregulated genes are hypermethylated when compared to upregulated genes, however no clear differences were observed between both conditions. C) Violin plot showing distribution of methylation at TSS of genes classified into quartiles according to expression level (highest, 4). Each violin is scaled to the same maximum width (total area is not constant between violins) to demonstrate distributions for each quartile. Black dots denote the median.

Six genes were downregulated by more than 4-fold in offspring of hypoxia exposed males (Figure 4), including *timm23b* (translocase of inner mitochondrial membrane 23 homolog b (yeast)), *acot21* (acyl-CoA thioesterase 21), *znf1156* (zinc finger protein 1156), *cyt1l* (type I cytokeratin), *irge1* (immunity-related GTP-ase), and *zgc:158417*.

Of the genes we analysed, 13,698 were associated with a GO term (including 58 of the 91 differentially expressed genes). Nine GO terms were overrepresented in our differentially expressed genes at a q-value cutoff of 0.1 (Table 1). Several significantly enriched GO terms were associated with lytic or proteolytic activity (i.e. serine hydrolase activity, serine-type endopeptidase activity, hydrolase activity, serine-type peptidase activity, glutamate decarboxylase activity, endopeptidase activity, peptidase activity, peptidase activity, acting on L-amino acid peptides). Proteolysis (GO:0006508 p-value = 7.43 x 10^−04^; q-value = 0.7640) tended to be over-represented. Genes associated with the process of aging (GO:0007568; p-value = 1.99 x 10^−05^; q-value = 0.1640) as well as to the response to oxidative stress (GO:0006979; p-value = 9.20 x10^−04^; q-value = 0.841) also tended to be over-represented.

**Table 1:** Gene Ontology analysis terms associated with differentially expressed genes. Only terms with a False Discovery Rate q-value below 0.1 are displayed.

| GO term | Sub-ontology | Description | Enrichment factor | p-value | FDR q-value |
| --- | --- | --- | --- | --- | --- |
| GO:0004252 | function | serine-type endopeptidase activity | 15.97 | $2.47 \times 10^{-07}$ | $7.55 \times 10^{-04}$ |
| GO:0008236 | function | serine-type peptidase activity | 13.67 | $7.18 \times 10^{-07}$ | $1.10 \times 10^{-03}$ |
| GO:0017171 | function | serine hydrolase activity | 13.67 | $7.18 \times 10^{-07}$ | $7.30 \times 10^{-04}$ |
| GO:0016787 | function | hydrolase activity | 2.62 | $1.56 \times 10^{-05}$ | $1.19 \times 10^{-02}$ |
| GO:0004351 | function | glutamate decarboxylase activity | 216.71 | $2.09 \times 10^{-05}$ | $1.28 \times 10^{-02}$ |
| GO:0004175 | function | endopeptidase activity | 6.35 | $3.34 \times 10^{-05}$ | $1.70 \times 10^{-02}$ |
| GO:0070011 | function | peptidase activity, acting on L-amino acid peptides | 4.84 | $8.66 \times 10^{-05}$ | $3.77 \times 10^{-02}$ |
| GO:0008233 | function | peptidase activity | 4.64 | $1.19 \times 10^{-04}$ | $4.53 \times 10^{-02}$ |
| GO:0009449 | process | gamma-aminobutyric acid biosynthetic process | 216.71 | $2.09 \times 10^{-05}$ | $1.28 \times 10^{-02}$ |

### Global methylation changes in the sperm of parental males and the relationship between methylation in parental sperm and gene expression in offspring

To explore potential mechanistic explanations underlying increased tolerance to acute hypoxia and differential gene expression in the zebrafish offspring, we conducted whole genome bisulfite sequencing (WGBS) of sperm from hypoxic (*n*= 3) and control males (*n*=3). We obtained an average of 67,891,334 reads per methylome (SEM ± 2,551,964) and a mapping efficiency of 45.02% (SEM ± 0.07%). Global cytosine-guanine (CG) dinucleotide methylation levels showed no differences between hypoxia and control samples (84.15% and 84.23%, respectively; *p* > 0.05).

In jawed vertebrates, methylation of transcription start sites (TSS), and in particular regions enriched on CG dinucleotides (CGIs), is associated with transcriptional silencing (Peat et al., 2014). For two example cases, the upregulated hemoglobin cluster and the downregulated *acot21* gene, we characterized methylation at individual positions and CGI using statistical and experimental criteria (Gardiner-Garden and Frommer, 1987; Long et al., 2013). No differences in methylation levels were observed for these genes (Figure 5A). Next, we explored methylation levels for all differentially expressed genes. Using a threshold of 20 methylation calls we obtained 29 and 23 genes for the upregulated and downregulated groups. Whereas downregulated genes are hypermethylated and upregulated genes are hypomethylated for hypoxia and control samples, TSS methylation values remain stable when both groups of samples are compared (Figure 5B). Finally, we explored the global effect of DNA methylation on gene expression using expression quantiles. We found methylation levels in the parental sperm and gene expression in the F1 offspring are coupled in hypoxia and control samples (Figure 5C).

## DISCUSSION

We demonstrate that paternal exposure to hypoxia alters both the phenotypic response to hypoxia and gene expression in the offspring. Larvae of fathers that experienced moderate hypoxia-maintained equilibrium in acute hypoxia for longer than those of controls, indicating that paternal exposure stimulated a higher tolerance to hypoxic conditions. Using next-generation sequencing, we also detected significant changes in gene expression between control and hypoxia offspring, with two key hemoglobin genes upregulated in offspring of hypoxia exposed males −genes which may mediate the observed phenotypic differences, as they are involved in oxygen transport. This pattern of inheritance, through the paternal line, could have large evolutionary consequences as fathers are able to pass down valuable information to offspring that may enable better survival. However, the underlying mechanism for this transmission remains unknown, as we did not detect any differential methylation in the sperm of parental males at any of the differentially expressed genes.

Importantly, our study provides evidence of increased tolerance to acute hypoxia through paternal exposure, thus, adding to increasing evidence that environmental challenges experienced by ancestors can provide progeny with environmental specific information that might allow future generations to survive the same environmental challenge, i.e., transgenerational plasticity (Donelson et al., 2014, 2012; Heckwolf et al., 2018; Herman and Sultan, 2011; Lee et al., 2020; Marshall, 2008; Ryu et al., 2018; Veilleux et al., 2015). In a previous study, 20 dpf larvae exposed to 4 kPa pO_2_ revealed that offspring of parents exposed to hypoxia had longer time to loss of equilibrium (hypoxia resistance; (Ho and Burggren, 2012)). In preliminary trials, we found no behavioral differences using these oxygen parameters; thus, we increased the magnitude of hypoxia, exposing larvae to ~1 kPa, which produced the expected phenotypic differences. Importantly, the previous study (Ho and Burggren, 2012) did not differentiate between maternal and paternal effects and did not provide a possible mechanism underlying this phenotypic effect.

It is important to note that our study detects intergenerational acclimation, with potential for transgenerational acclimation. A transgenerational study requires rearing fish through to create an F2 generation. Indeed, few studies that claim transgenerational acclimation are truly transgenerational, as the studies are conducted across a single generation (O’Dea et al., 2016). Even if parents are exposed to environmental challenges prior to maturity, the primordial germ cells are still exposed to the challenge as well. Regardless, our results suggest that parents are passing on information that may benefit offspring survival, thus facilitating acclimation to environmental conditions.

To try and understand the underlying mechanism priming progeny to better cope with hypoxic conditions, we conducted transcriptomic analysis of the progeny. We detected 91 genes that were differentially expressed in the offspring of paternal males that were exposed to moderate hypoxia for two-weeks. Most notably, two hemoglobin genes (*hbaa1* and *hbz*) exhibited over 7-fold differential expression, were upregulated in offspring of males exposed to hypoxia, with another hemoglobin gene, *hbba1*, also upregulated, though non-significantly. Remarkably, sequenced offspring from the H1 and H2 families, the two families that show the greatest tolerance to acute hypoxia also show the greatest upregulation in hemoglobin gene expression (Table S3). *hbaa1, hbz* and *hbba1* are all found on chromosome 3, and are part of the major hemoglobin locus responsible for heme binding, and are instrumental in oxygen transport (Ganis et al., 2012). Exposing fish to hypoxia is typically considered to improve hypoxia tolerance through alterations in hemoglobin, hemoglobin-O_2_ binding affinities, or cardiac function to improve low O_2_ performance (Cook et al., 2013). Thus, differential expression of hemoglobin genes (up-regulated in offspring from hypoxia exposed fathers) could indicate altered physiological mechanisms to combat low oxygen, which could be precipitating the increased tolerance to acute hypoxia in the H1 and H2 families.

The relationship between hypoxia and hemoglobin function has been studied in larval zebrafish, suggesting that zebrafish larvae might be able to upregulate hemoglobin concentration in response to chronic hypoxia (Schwerte et al., 2003). Under normoxic conditions, oxygen supply via diffusion seems to be sufficient to meet metabolic demands up to 12-14 dpf in zebrafish, but larvae appear to be able to use a circulatory system as a backup where necessary (Schwerte et al., 2003). Further, impaired hemoglobin function does not impair routine oxygen usage in normoxia or at moderate levels of hypoxia in 5-42 dpf larvae, but functional hemoglobin does allow larvae to sequester extra oxygen from water in extreme hypoxic conditions (Rombough and Drader, 2009). Larvae express embryonic/larval globins early in development, but somewhere between days 16 and 22 the embryonic/larval globins begin to decline (Ganis et al., 2012; Tiedke et al., 2011), and adult globin expression increases, with the adult globin expression pattern nearly completely established by day 32dpf. *hbae5* (ENSDARG00000045142) is the only embryonic/larval hemoglobin to show any pattern of differential expression in our transcriptomic data. We found that this gene was highly expressed and upregulated twice as much in our hypoxia offspring than in our control, though not significantly differentiated after FDR correction (p= 0.014, q = 0.391). *hbae5* expression appears to peak around 22 dpf (Ganis et al., 2012), so upregulation of this gene should result in higher affinity for oxygen, which may help explain the increased tolerance to the acute hypoxia that we observed in our 20-21 dpf progeny. *hbae5* has also been shown to be up-regulated more than 5-fold by hypoxia in zebrafish larvae directly exposed to hypoxia; ((Long et al., 2015); note that hbae5 is called hbz in their study).

In addition to two genes upregulated over 7-fold in offspring of hypoxia exposed males, six genes were downregulated by more than 4-fold as well. These include *timm23b* (translocase of inner mitochondrial membrane 23 homolog b (yeast)), an integral component of membranes. Gene *acot21* (acyl-CoA thioesterase 12), is a key component of acyl-CoA metabolic processes with thiolester hydrolase activity and found in the cytoplasm. Gene *znf1156* (zinc finger protein 1156) has metal ion binding functions. Gene *zgc:158417* and *irge1* (immunity-related GTP-ase family) are predicted to be integral component of membranes and used in GTP binding. Gene *cyt1l* (type I cytokeratin) is predicted to have structural molecule activity. While there is little information on these downregulated genes, expression changes of such significant magnitude are likely to have large consequences.

An HRGFish database (Rashid et al., 2017) reports 50 key genes that are altered through hypoxia, but we did not find any of them to be differentially expressed in our transcriptomic data (Table S1, showing data for 46 of the 50 HRGFish genes referenced for the zebrafish), suggesting that the genes that are altered by direct exposure to hypoxia may be quite different to those that may be passed on to future generations of hypoxia exposed ancestors. Importantly, transcriptomic studies of F1 and F2 adult tissues, like heart, gill, brain and liver, might reveal differential expression of some of these key hypoxia genes.

Interestingly, our GO analysis highlighted an overrepresentation of genes associated with aging and oxidative stress. Studies have previously hypothesized a link between hypoxia (and effects on respiration), oxidative stress (and free radical production) and aging in humans (Katschinski, 2006; Valli et al., 2015), with several studies demonstrating age-related changes in the hypoxia inducible factor system.

DNA methylation is known to be altered by hypoxia exposure in fishes (Lai et al., 2019; Wang et al., 2016) and other vertebrates (Childebayeva et al., 2019; Zhang et al., 2017). Thus, we hypothesized that hypoxia might produce changes in the paternal methylome, which are inherited to the offspring, explaining the observed tolerance to acute hypoxia and differential gene expression. Interesting, we found methylation levels and gene expression were coupled in our hypoxia and control samples, despite the different tissues of origin for the methylome and RNA expression data. This coupling supports a common pattern between the paternal methylome and expression levels of the embryo. Yet, we observed no differences in methylation levels in any differentially expressed genes, including the hemoglobin cluster, suggesting that sperm methylation patterns are not responsible for changes in the observed changes in gene expression in our study, or perhaps through epigenetic control away from the focal genes. The medaka studies (Lai et al., 2019; Wang et al., 2016) exposed fish to a similar level of hypoxia, but for a much longer period, the entire 3-month life cycle, which may explain why they observed differential methylation in the sperm and we did not. If alterations to DNA methylation are not responsible for the changes in gene expression observed in our study, then perhaps histone modifications or miRNAs are at play. miRNA in medaka have also been identified as a potential underlying mechanism of transgenerational testis impairment induced by hypoxia by targeting genes associated with stress responses, cell cycle, epigenetic modifications, sugar metabolism, and cell motion (Tse et al., 2016).

In conclusion, our study demonstrates that paternal exposure to hypoxia is able to endow offspring with a higher resistance to hypoxic conditions and that upregulation of genes at the hemoglobin gene complex on chromosome 3 may explain these phenotypic effects. This study is the first to demonstrate that paternal exposure alone can mediate these changes in phenotype and gene expression. These findings suggest that paternal inheritance may be providing offspring with valuable environmental specific information (a ‘memory’) that could increase their survival, though it is unclear how this information is transmitted as we did not detect any differential methylation in the sperm of control and hypoxia treated males. Regardless, the variability in tolerance to acute hypoxia that we observed between hypoxia families suggests that progeny of different genotypes respond to hypoxia in different ways, phenotypically (i.e., an epigenotype x environment interaction). In other words, environmental specific information is transferred through some genotypes, but not others, suggesting that pre-acclimation to hypoxic conditions may be driven by an interaction between the genotype, the epigenotype and the environment.

## Acknowledgements

We thank Denham Cook for advice on designing our hypoxia system, Murray McKenzie for construction of the hypoxia system, and Sean Divers for help with set up and construction of the acute hypoxia assay chambers. Greg Gimenez provided advice on RNASeq analyses and Noel Jhinku and other members of the Otago Zebrafish Facility assisted with fish husbandry.

## Funding

This work was supported by a Royal Society of New Zealand Marsden Fund grant (UOO1507) and a University of Otago Research Grant to S.L.J. The funders had no role in study design, data collection and analysis, decision to publish, or preparation of the manuscript.

## Authors’ contributions

S.L.J., T.H. and N.G. conceived the study. A.R., A.B., J.C., and T.K. collected the data. A.R., S.L.J., O.O. and L.D. performed the analyses. A.R. and S.L.J led the writing of the manuscript, with assistance from O.O. and L.D. for bioinformatics aspects, and all the authors contributed to revisions.

## Competing interests

The authors declare no competing financial interests.

## Data and materials accessibility

The raw phenotypic data can be found at https://osf.io/xpy8j/. The accession number for the RNA-Seq and WGBS datasets reported in this paper is GEO:GSE160662. The source code for the bioinformatic analysis is publicly available on GitHub https://github.com/OscarOrt/Paternal_hypoxia_Ragsdale_2020.

## SUPPLEMENTARY MATERIALS

Supplementary material for this article is available at…

Table S1. Results of differential gene expression analysis

Table S2. Read counts mapping to four hemoglobin genes

Table S3. Whole genome bisulfite sequencing summary data

---

**Table S1.**
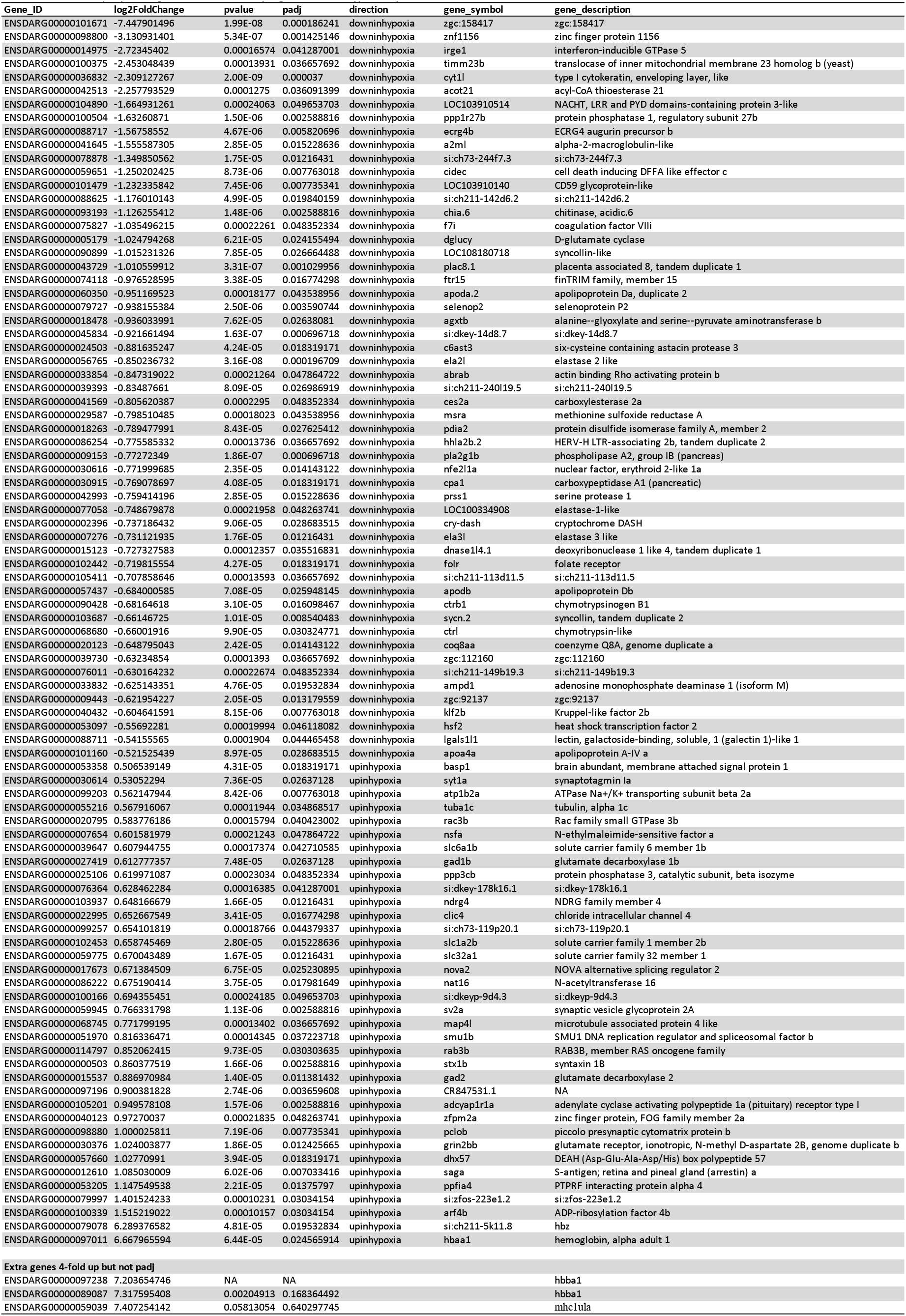
Differentially expressed genes between the offspring of control vs. hypoxia exposed males.

**Table S2:** Read counts mapping to four haemoglobin genes located on a small region of chromosome 3. * genes are not significantly differentially expressed but show strong expression in samples H1 and H2 who were the most resilient to loss of equilibrium in hypoxic environment. Note that there are two versions of hbba1 due to genome duplication, and they are physically separated on chromosome 3.

|  | Gene name | C1 | C2 | C3 | H1 | H2 | H3 |
| --- | --- | --- | --- | --- | --- | --- | --- |
| **Ensembl ID** |  |  |  |  |  |  |  |
| **ENSDARG00000079078** | *hbz* | 0 | 0 | 2 | 9 | 303 | 1 |
| **ENSDARG00000097011** | *hbba1* | 2 | 1 | 17 | 671 | 1427 | 14 |
| **ENSDARG00000097238** | *hbba1\** | 0 | 0 | 8 | 341 | 974 | 8 |
| **ENSDARG00000089087** | *hbba1\** | 1 | 3 | 13 | 227 | 1169 | 21 |

**Table S3.**
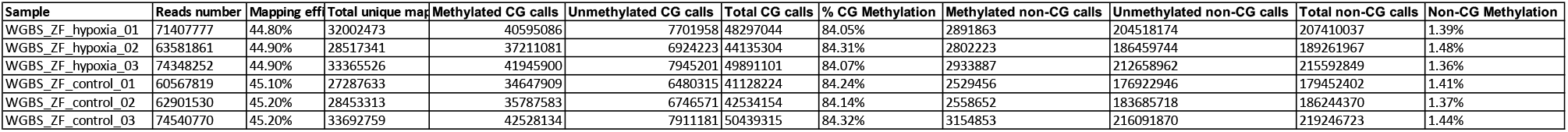
Whole genome bisulfite sequencing of the sperm of hypoxic vs. control males. The table lists the general sequencing statistics as well as the number of cytosine calls at either CG dinucleotides (‘CG’) or in other sequence contexts (‘non-CG’), for the samples used in the experiments, mapped against the Zebrafish genome version 11 (GRCz11). Details of bioinformatic processing are provided in the Methods section. The frequency of non-CG methylation indicates the maximum rate of non-conversion during the bisulfite treatment step; by this measure, all libraries had a bisulfite conversion efficiency of at least 98.52%.

## Notes

https://osf.io/xpy8j/

https://github.com/OscarOrt/Paternal_hypoxia_Ragsdale_2020

### Competing Interest Statement

The authors have declared no competing interest.

